# FlywheelTools: Data Curation and Manipulation on the Flywheel Platform

**DOI:** 10.1101/2021.03.12.434998

**Authors:** Tinashe M. Tapera, Matthew Cieslak, Max Bertolero, Azeez Adebimpe, Geoffrey K. Aguirre, Ellyn R. Butler, Philip A. Cook, Diego Davila, Mark A. Elliott, Sophia Linguiti, Kristin Murtha, William Tackett, John A. Detre, Theodore D. Satterthwaite

## Abstract

The recent and growing focus on reproducibility in neuroimaging studies has led many major academic centers to use cloud-based imaging databases for storing, analyzing, and sharing complex imaging data. Flywheel is one such database platform that offers easily accessible, large-scale data management, along with a framework for reproducible analyses through containerized pipelines. The Brain Imaging Data Structure (BIDS) is a data storage specification for neuroimaging data, but curating neuroimaging data into BIDS can be a challenging and time-consuming task. In particular, standard solutions for BIDS curation are not designed for use on cloud-based systems such as Flywheel. To address these challenges, we developed “FlywheelTools”, a software toolbox for reproducible data curation and manipulation on Flywheel. FlywheelTools includes two elements: *fw-heudiconv*, for heuristic-driven curation of data into BIDS, and *flaudit*, which audits and inventories projects on Flywheel. Together, these tools accelerate reproducible neuroscience research on the widely used Flywheel platform.

## 1 INTRODUCTION

Many fields in science are grappling with failures of scientific reproducibility (Botvinik-Nezer et al. 2020). Given the high dimensionality of the data, the need for complex image processing, and a plethora of analytic techniques, this crisis is particularly acute for neuroimaging research. Increasingly, the field has recognized that unstructured filesystems are a major source of non-reproducible practices. As such, major academic centers and large consortia have increasingly adopted platforms that leverage database technologies that have become standard in other fields. In addition to providing functionality for searching and categorizing complex source data, imaging databases enhance reproducible research by providing a clear audit trail of image processing applied to the data and its results, including both derived images and other data. Widely used imaging databases include Collaborative Informatics Neuroimaging Suite (COINS) (Landis et al. 2016), eXtensible Neuroimaging Archive Toolkit (XNAT) (Herrick et al. 2016), Longitudinal Online Research and Imaging System (LORIS) (Vaccarino et al. 2018), and others (Book et al. 2016; Helmer et al. 2011; Rogovin et al. 2020; Helmer et al. 2011; Poldrack and Gorgolewski 2017). More recently, the commercial platform Flywheel has seen growing adoption by major academic imaging centers (e.g., Stanford, Columbia, Duke, and Penn) due to its modern technology, ease of use, and scalability.

Most neuroimaging databases now leverage the standards defined by the Brain Imaging Data Structure (BIDS) (Gorgolewski et al. 2016). BIDS is an open-source standard for neuroimaging data organization that specifies how files should be named, how directories should be organized, and how metadata should be structured. As such, BIDS provides users with a well-documented format to understand both imaging data and metadata. Importantly, as BIDS provides a transparent format for recording imaging parameters and key aspects of the experimental design, BIDS enhances both data accessibility and data sharing.

Furthermore, BIDS allows users to leverage BIDS-apps — image processing pipelines, such as fMRIPrep, c-Pac, and QSIprep, that read the metadata defined by BIDS (Esteban et al. 2019; Craddock et al. 2013; Cieslak et al. 2020). As BIDS-apps can auto-configure to ensure that analytic parameters are appropriate for the input data provided, they dramatically reduce barriers to implementing best practices in image processing. Furthermore, containerized BIDS-apps encompass all software dependencies, further enhancing reproducibility.

On a filesystem, conversion of raw DICOM images to NIfTIs that conform to BIDS can be accomplished with a variety of tools including *HeuDiConv, dcm2bids*, and others. However, this crucial step, a process typically called “BIDS curation,” is often difficult to implement within the database environment, as it requires a mechanism for converting data in hierarchical databases into BIDS. As BIDS curation is one of the very first steps performed on the data, lack of transparency in curation can impact the reproducibility of all subsequent steps. Here we introduce FlywheelTools, a software suite that provides flexible and reproducible methods for BIDS curation on the Flywheel platform. Documentation and code presented in this work can be found online at: https://fw-heudiconv.readthedocs.io/en/latest/.

## 2 METHODS

The FlywheelTools toolkit allows users to follow a reproducible workflow for BIDS curation and auditing of their data. This workflow typically includes: inspection of sequences collected during a study, design of a curation schema, implementation of that curation schema, and, finally, auditing and inspection of the curated data.

### 2.1 Programming Languages & Technologies

FlywheelTools is built primarily in Python 3.6 (Van Rossum and Drake 2009) to leverage Flywheel’s highly accessible Software Development Kit (SDK). Additionally, R 3.4.1 (R Core Team 2019) is used for HTML report generation. For reproducibility and workflow management, the modules of FlywheelTools are packaged in version-controlled software containers built and managed in Docker (Merkel 2014). Lastly, the FlywheelTools package relies on users adopting BIDS to curate their data. BIDS has rapidly evolved to become the directory standard in the neuroimaging community for reproducible data organization. Importantly, BIDS is supported by a large community that contributes to its development and adoption. Further, proposals for BIDS schema pass through a rigorous testing process before being adopted. Software developers leverage this ubiquity by creating BIDS-apps: analysis and processing pipelines that operate directly on BIDS datasets as inputs, enhancing reproducibility and interoperability.

### 2.2 Flywheel

Flywheel is a data management and analysis platform that is tailored for neuroimaging research. The platform focuses heavily on collaborative and reproducible science. User-facing components of the platform itself are the web User Interface (UI), the Command Line Interface (CLI), the Flywheel Software Development Kit (SDK), and the Application Programming Interface (API).

### 2.3 Flywheel Web UI

The web UI is accessible through any modern web browser. Through this point-and-click interface, users are able to upload, view, download, and analyze data with ease. However, accomplishing tasks with many repetitive steps or over a large number of participants/sessions can be tiresome and error-prone. Alongside knowledge of navigating the web UI, many users also make use of the API and SDK to manipulate and analyze data programmatically.

### 2.4 Flywheel API & SDK

Flywheel’s database uses MongoDB for data storage and access, meaning that all Flywheel data are represented by hierarchical relationships between document objects. This allows users to create and store complex structures with ease, and query data rapidly (Banker 2011). To access these data, Flywheel uses a RESTful Application Programming Interface (REpresentational State Transfer) (Biehl 2016), making each document or data object accessible through a specific URL that a web browser or SDK can access by requesting the data and waiting for a response from the server. The Flywheel Python SDK^1^ provides a powerful interface for inspecting and manipulating data through this API. By standardising this underlying data model into Pythonic objects, the Flywheel SDK is effectively an object relationship mapper, similar to the popular SQLAlchemy software.

### 2.5 Flywheel Data Model

Objects in Flywheel’s data model follow a specific hierarchical structure — at the top level is a Flywheel instance, a process that serves the API to an organization (for example, a neuroimaging center). Within the Flywheel instance, there are multiple groups, which are typically labs or research units that collaborate on one or more projects. Each project object can have one or many subjects (i.e. participants), and each subject can have one or many sessions (i.e. scanning visits). Within a session, there may be one or many acquisition objects which represent the scanning sequences collected during a particular scan or examination (e.g., sMRI, rs-fMRI, dMRI). Finally, the data files associated with the sequence (e.g., NIfTIs or DICOMs) are attached to each acquisition. Note that a file can additionally be attached to any object type, and each object can have metadata associated with it. Hence, a “subject” object may have metadata associated with that participant (such as demographic information) and may also have a text file attached to it (such as clinical data). A notable exception to this hierarchical structure is the analysis object, which behaves in much the same way as others but can be a child object of any other object, allowing researchers to create analyses of entire projects, for example, each with their own associated metadata and files.

Abstracting this data model in Python results in simple hierarchical objects, each with methods for handling metadata and files, and methods for accomplishing object-specific tasks like traversing the hierarchical structure or running analyses. The modules of FlywheelTools make use of this data model to accomplish a wide range of tasks.

### 2.6 Flywheel Gears

Flywheel encourages the use of pre-packaged computational workflows, called “gears”. Gears are run by virtual machines/containers using Docker and hence are version-controlled and software/platform agnostic. Gears can accomplish tasks such as data manipulation, pre-processing, and analysis. In addition to the existing gears available on the platform, users are able to package their own software in a gear and use it for running analysis workflows on their Flywheel data via the web UI or SDK. The complexity and frequency of the task guides if a task should be accomplished using the web UI, programmatically using the SDK, or by wrapping it as a workflow into a gear. Gears ingest existing Flywheel data (such as images or file attachments) as inputs to the workflow and can be created with clickable configuration options. Once a workflow has completed running, Flywheel collects any files remaining in the pre-defined output directory of the container and attaches them to a resulting analysis object. The output of a gear (such as an HTML report or tabulated data) can be viewed on the Flywheel UI, downloaded to disk for further sharing or analysis, or used as input to a subsequent gear.

## 3 RESULTS

FlywheelTools is implemented using the Flywheel SDK to enable easy inspection, curation, validation, and audit of Flywheel data through a handful of user-friendly gears and command-line interfaces. The first module of the package is called *fw-heudiconv* and is largely inspired by the popular Heuristic DICOM Converter package (Halchenko et al. 2018). *fw-heudiconv* is a multi-part toolbox for reproducible curation of neuroimaging data into BIDS on Flywheel. The second module, *flaudit*, is a tool for auditing a Flywheel project, giving users an overview of the key elements of their data set.

### 3.1 FW-HEUDICONV

The first tool, *fw-heudiconv*, is a multi-purpose command-line interface and Flywheel gear designed for BIDS curation on Flywheel (**Figure 1**). It is designed to be intuitive, flexible, and reproducible.

**Figure 1:**
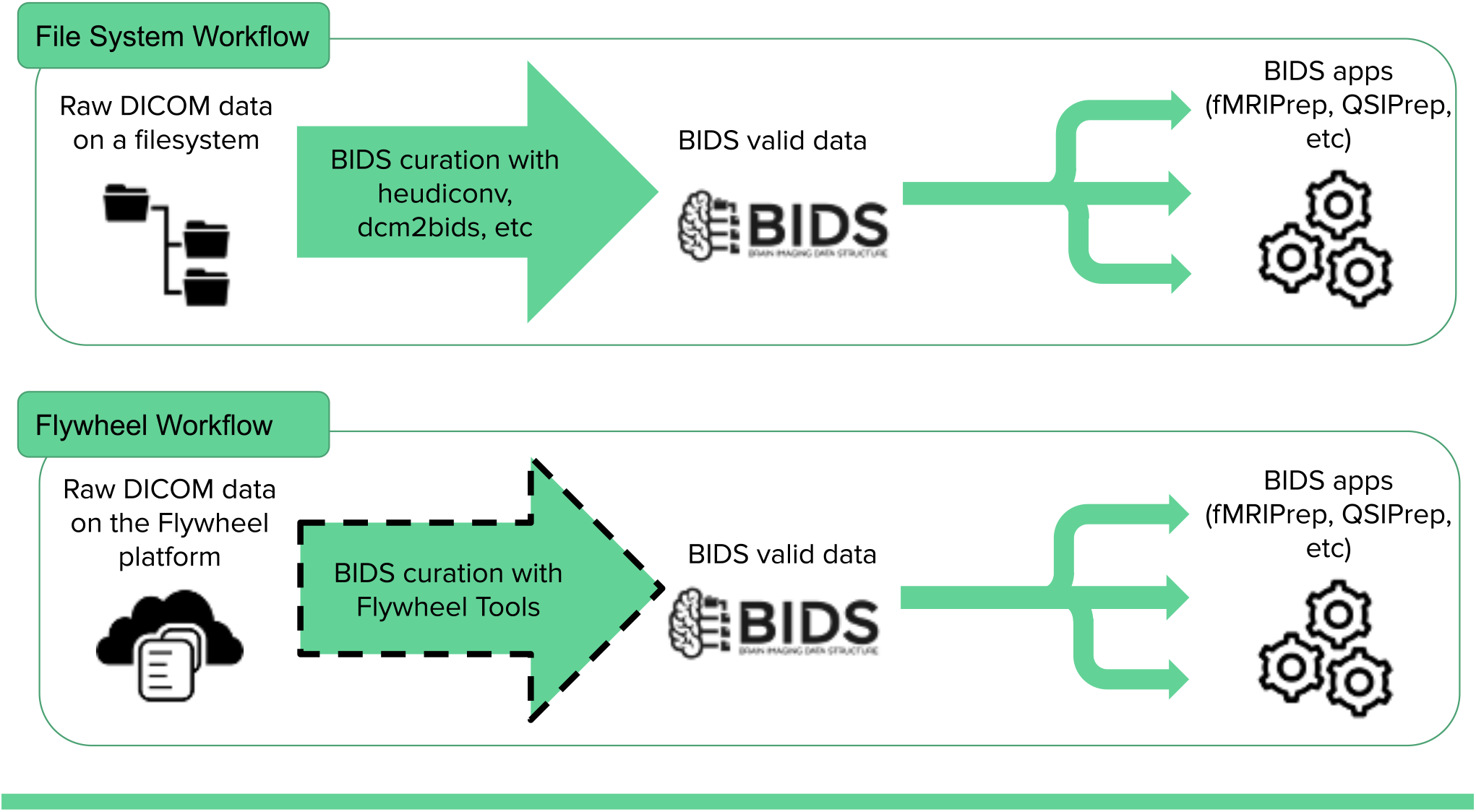
BIDS-app workflow. BIDS curation on file systems is a common task that can be accomplished by existing tools (heudiconv, dcm2bids, etc) or manually, but mechanisms for BIDS curation on many cloud databases have yet to be developed. FlywheelTools provides this functionality for the Flywheel platform.

#### 3.1.1 Architecture & Design

*fw-heudiconv* is inspired in large part by the Heuristic DICOM Converter (*HeuDiConv*) package, and shares much of its design practices. To curate data into BIDS format, *fw-heudiconv* first considers DICOM data to be the “ground truth” and builds its curation approach using data in the DICOM headers. On Flywheel, when DICOMs are added to the database, either through manual upload or automatically from a linked scanner. The DICOMs are then automatically converted into NIfTI files by Flywheel’s automated gears. The result is an acquisition object with both DICOMs and one or more NIfTIs. Ultimately, *fw-heudiconv* only has permission to manipulate metadata associated with a NIfTI file. By not manipulating DICOMs or their associated metadata, BIDS curation can safely be reproduced from ground truth data.

*fw-heudiconv* can be downloaded as a Python command-line interface from the Python Package Index using *pip*. It is also available as a point-and-click gear that is managed by Docker containerization. This containerization allows for reproducible version control, as every computation can be recorded, including information about the software, OS, and algorithm versions. There are a number of commands available in *fw-heudiconv*, and each of them starts by querying data from Flywheel. Users can filter their queries to operate on an entire Flywheel project, a subset of subjects, or a subset of sessions. Notably, with the `--dry-run` option, each command has the ability to test and evaluate its effects without actually manipulating metadata in the Flywheel database or writing data to disk. Below, we consider each of the five available commands.

##### 3.1.1.1 fw-heudiconv-tabulate

The tabulate tool is used to parse and extract DICOM header information in a project (or within a filtered subset of that project) and compile these data into a table for the user to examine. By collecting DICOM header information into a tabular format, the tabulate tool gives users a comprehensive overview of the different scanning sequences that have been collected in the query, including the sequence parameters. Additionally, users have the option to limit the tabulation to a unique combination of common DICOM header fields, which significantly decreases the complexity of the table. When used at the command line, the table produced by this command is written to a local disk. As a gear on Flywheel, the table is automatically saved in the output section of the gear.

##### 3.1.1.2 fw-heudiconv-curate

The curate tool is used to curate a dataset on Flywheel into BIDS format. Much like *HeuDiConv*, curation is accomplished through the use of a heuristic: a Python file that programmatically defines the templates for a range of BIDS-valid filenames, and defines the boolean logic that would assign a given scanning sequence to each template. This boolean logic is usually based on the sequence information users find in the tabulation of sequences, but all fields available in the DICOM header can be used to determine which template a particular file can be assigned to. Additionally, the curate tool can be used to manipulate BIDS metadata that may need to be added to the dataset. The process of curation only manipulates the BIDS metadata of NIfTI files, and hence can be repeated or updated at any time at the user’s discretion.

##### 3.1.1.3 fw-heudiconv-export

The export tool is used to export a BIDS dataset on Flywheel to disk. It can also be used by other gears and scripts to quickly and easily extract their BIDS data into the workspace of their analysis pipeline.

##### 3.1.1.4 fw-heudiconv-validate

The validate tool is a wrapper around the popular BIDS Validator package and is used to check if the applied curation results in a BIDS-valid dataset. After exporting a dataset with *fw-heudiconv-export*, the validate tool runs the BIDS Validator on the dataset and returns the verbose output of the errors and warnings given by the BIDS Validator. Additionally, the results of the validator can be tabulated for easy inspection. On the Flywheel GUI, *fw-heudiconv-validate* also displays a green check mark in the analysis tab for a successful validation, and a red check mark otherwise, allowing for quick visual inspection of BIDS curation status for each session.

##### 3.1.1.5 fw-heudiconv-clear

The clear tool is used to clear BIDS information cleanly and safely from the project or subjects and sessions queried. This can be useful when a user wants to rerun the curation. The previously created persistent fields can be removed by running *fw-heudiconv-clear* before re-curating.

#### 3.1.2 The Heuristic File

The heuristic file is a Python file used as input to the*fw-heudiconv-curate* command. The file instructs *fw-heudiconv* on how to programmatically sort and parse through each acquisition object in Flywheel and assign it to a valid BIDS naming template. This is done by checking the attributes of a list of *seqInfo* objects — which are generated from each DICOM’s header information — against user-defined boolean rules. For example, if a T1-weighted image is present in a dataset, the user may define a string with a BIDS-valid naming template for this type of file, such as:

~~~
tlw = ‘sub-{SubjectLabel} ses-{SessionLabel} T1w.nii.gz’
~~~

Where the SubjectLabel and SessionLabel portions are expected to be automatically generated for each subject and session in the dataset. After the DICOM SeriesDescription field is added to the SeriesDescription attribute of *seqInfo*, the user can create a simple boolean expression to check if the string ‘T1w’ is in the SeriesDescription. If such a rule is met, this acquisition and its NIfTI file will be assigned to the T1-weighted image naming template. The NIfTI file will ultimately have this BIDS naming added to its metadata and be named correctly when exported to a filesystem. In more complex naming scenarios, *fw-heudiconv* can flexibly use boolean expressions involving any number of *seqInfo* attributes, which the user can access in the output of *fw-heudiconv-tabulate*.

In addition to setting naming templates, the heuristic file can also be used to hard-code and assign metadata in BIDS. These data are hard-coded into the metadata of the file object on Flywheel and are assigned by using specially reserved functions and keywords in *fw-heudiconv*. For example, the heuristic file can be used to point fieldmap scans to their intended sequences using a list:

~~~
IntendedFor = {
fieldmap1: [‘sub-{SubjectLabel}_ses-{SessionLabel}_task-rest_bold.nii.gz’]
}
~~~

By reserving select keywords for functions and metadata, heuristic files become versatile tools for defining and manipulating a wide array of metadata in Flywheel BIDS curation.

Importantly, because this heuristic file is plain text Python code, users are able to version control their files using Git and share these files via Github. Finally, when run on Flywheel as a gear, the heuristic file is automatically attached as an input to the analysis object created by *fw-heudiconv-curate*, allowing users to easily access the version history of their curation.

#### 3.1.3 Curation Workflow

For most users, the curation workflow follows the sequence detailed above (**Figure 2**). After DICOMs have been converted to NIfTIs, users can then begin by running *fw-heudiconv-tabulate* to gather the information stored in the DICOM headers necessary for creating a heuristic. Once the tabulation has been completed, the output file can be opened by any program that can read tabular data. At this stage, users can begin creating a heuristic file and running *fw-heudiconv-curate*, using the `--dry-run` flag to test the heuristic changes incrementally with informative logging. When satisfied, users can simply remove the `--dry-run` flag to apply the changes. The user can then use *fw-heudiconv-validate* to run the BIDS validator on the dataset or start over by removing all BIDS metadata with *fw-heudiconv-clear*.

**Figure 2:**
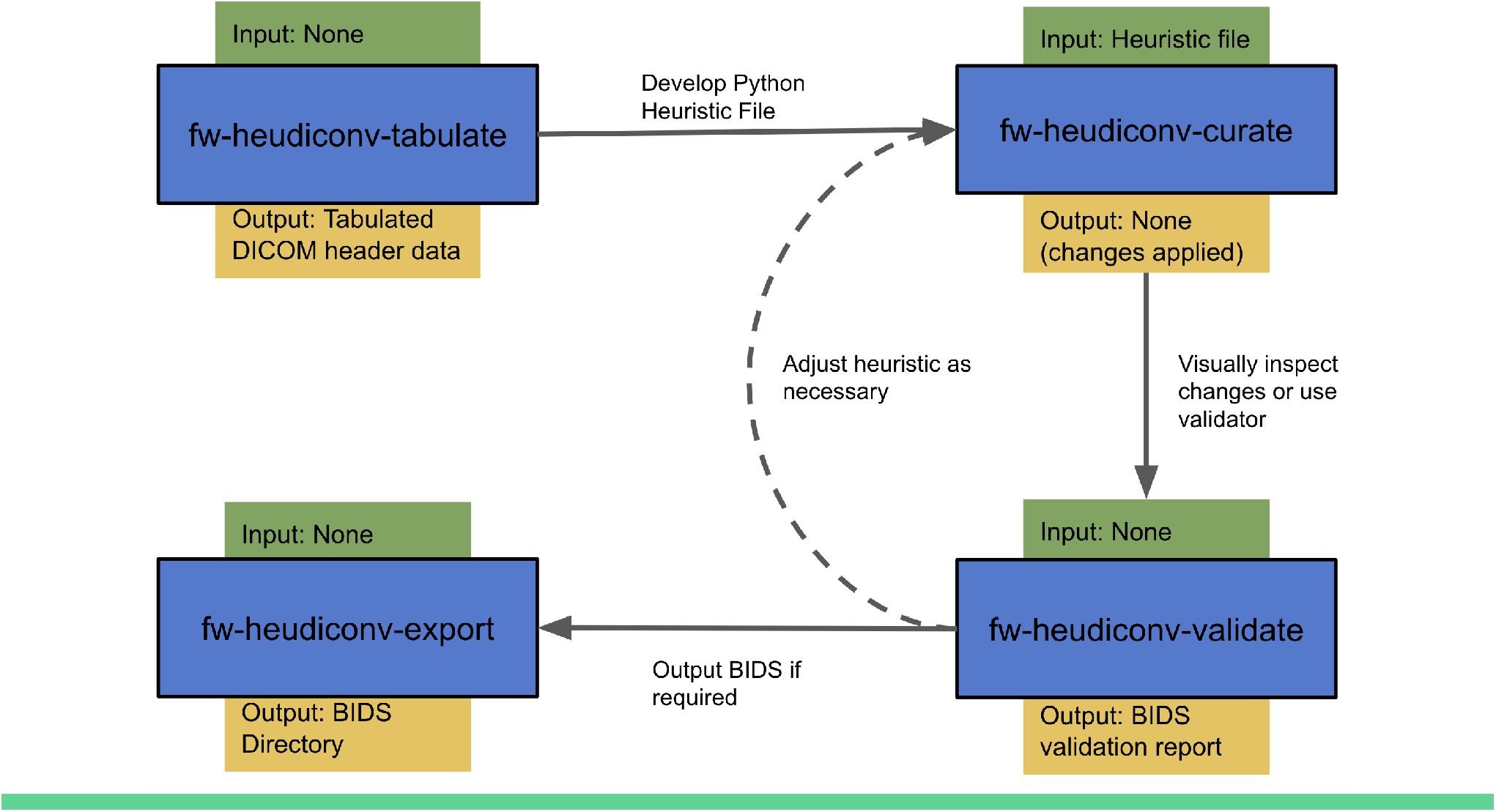
FlywheelTools workflow. Users first use the tabulate tool to extract sequence information from their data, which they use to develop a heuristic that delineates how sequences are mapped into BIDS. After this, they use the curate tool to convert their data into BIDS, and the validate tool to assess their curation. The export tool can be used to export their BIDS data as necessary.

If *fw-heudiconv* is run from the Flywheel GUI, each of the commands is available as a Flywheel gear. This option is beneficial for data provenance, as all of a gear’s commands and inputs, as well as outputs and logs, are stored and attached to each gear run.

### 3.2 FLAUDIT

The second module of FlywheelTools is a Flywheel project auditor, named *flaudit*. The module is intended to give Flywheel users a broad understanding of their entire Flywheel project, by summarizing the available data and illustrating analysis workflows. The output of this module, a portable HTML report, presents this information using a number of visualizations built in R Markdown using HTML, Javascript, and ggplot2, in two main sections: project overview and project completeness.

### 3.3 Architecture & Design

Using internal machinery similar to *fw-heudiconv-tabulate, flaudit* loops over existing data in a project and tabulates information about scanning sequences, BIDS metadata, and gear analyses that have been run. These three tables are saved internally and then passed as input to an R markdown script that generates an interactive HTML report. The data are also saved as output for the user to further access and analyze in their software of choice.

### 3.4 Flaudit: Project Overview

The overview section of the *flaudit* report provides a numerical overview of sequences, BIDS data, gear runs, and gear runtimes.

The first visualization uses the sequence data input to create a bar chart visualizing the names of the different sequences acquired across the entire Flywheel dataset. This visual is accompanied by an interactive table that users can search to compare values (**Figure 3**).

**Figure 3:**
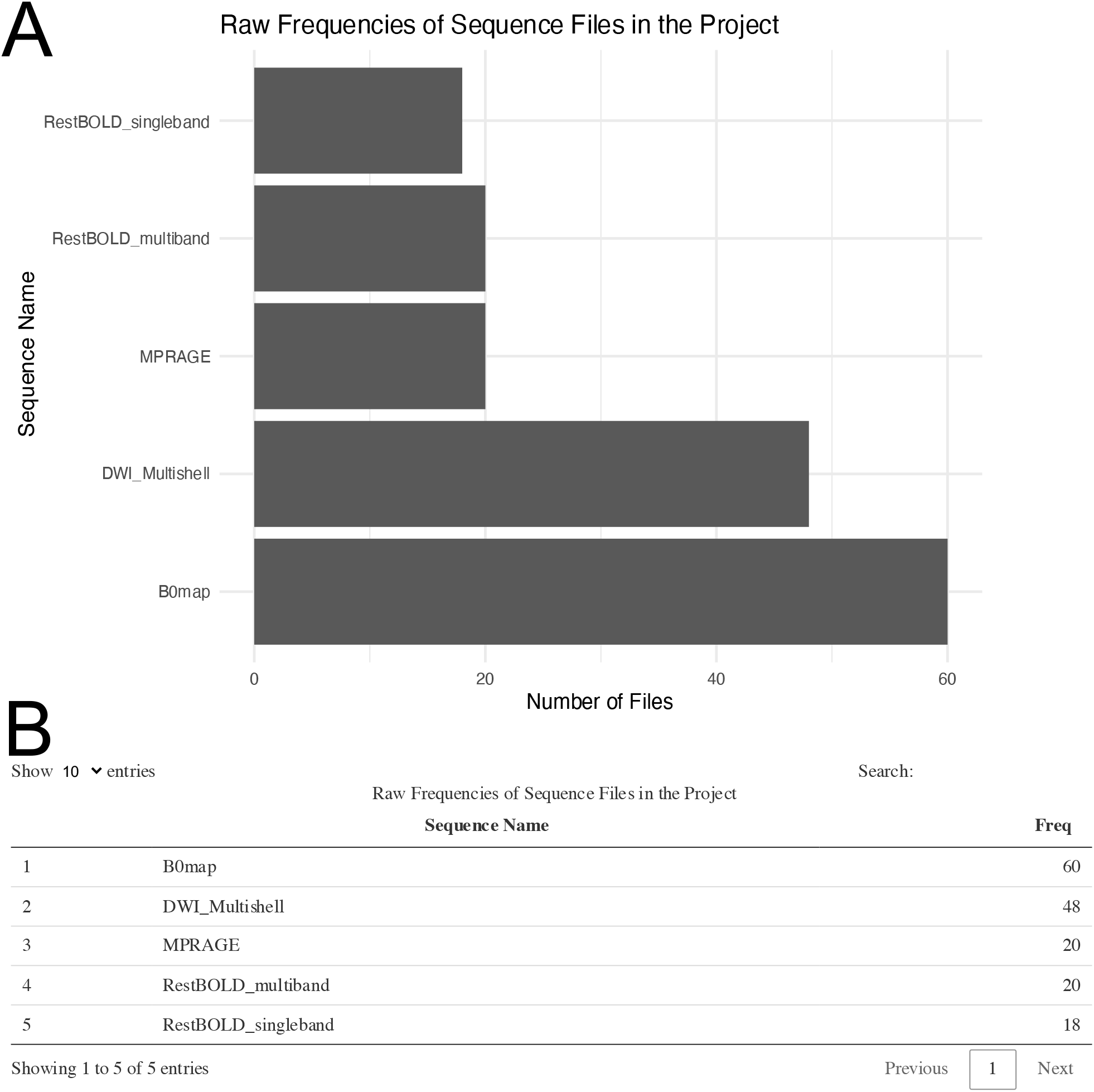
Enumeration of available sequences in a flywheel project. Panel (**A**) plots the count of files in each collected sequence; in this example, there are 60 files collected for the B0map sequence, as there are 20 subjects and 3 B0map sequences. Panel (**B**) shows the accompanying interactive table.

Next, using the BIDS metadata input, the report provides an interactive tree viewer to examine BIDS curation. In the tree, the nodes branch out from the project to show each sequence acquisition. For each acquisition, if the data has been curated into BIDS, the node itself can also branch out to show a BIDS name template, demonstrating what BIDS name that sequence has been given. Hovering over the BIDS name will display the number of subjects whose data have been named as such (**Figure 4**).

**Figure 4:**
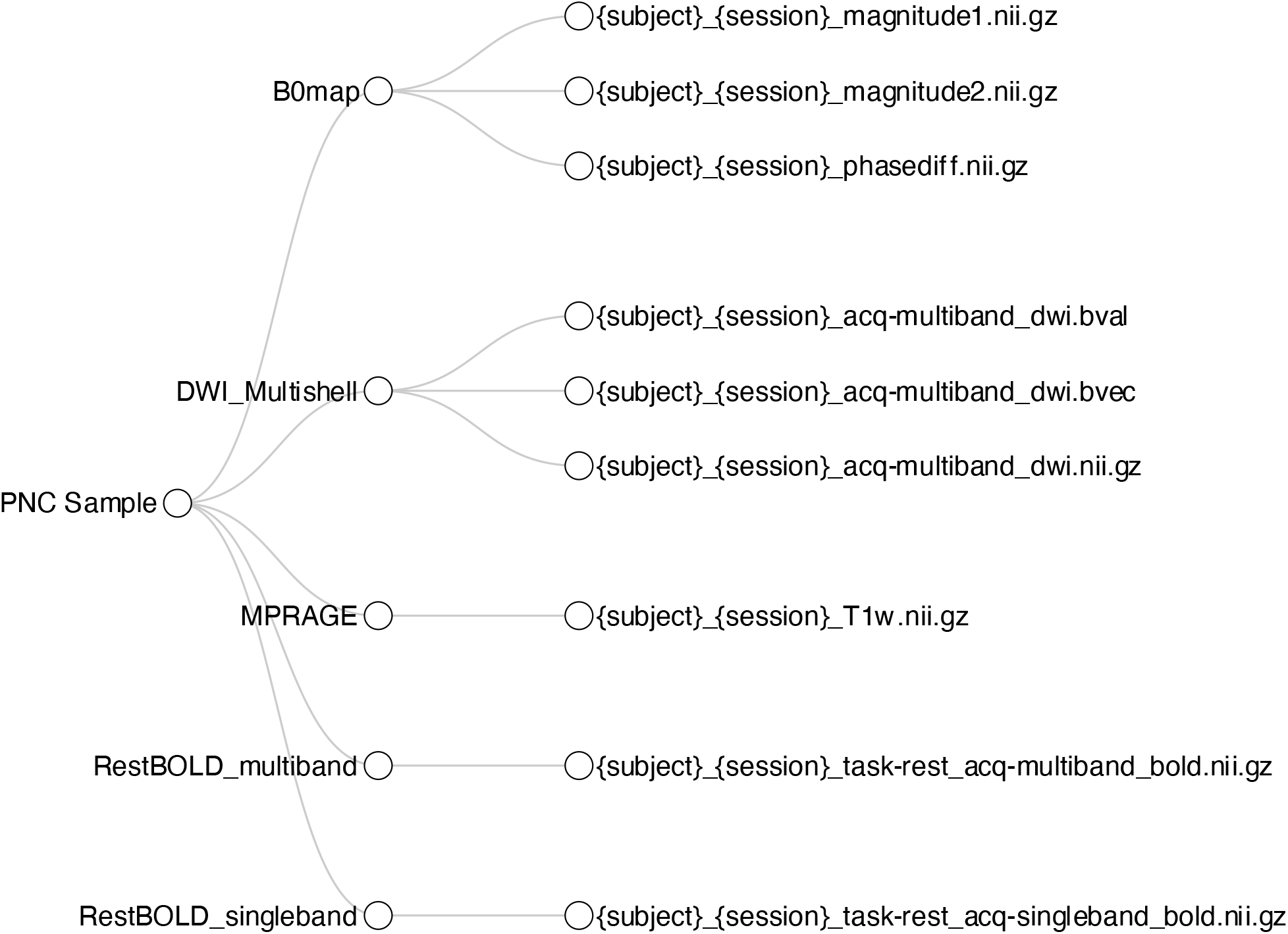
Interactive tree diagram illustrating BIDS curation. The tree shows how each sequence has been curated into BIDS format; users can hover their mouse over each leaf to show how many files have been curated into each BIDS filename template.

Finally, using the gear analysis data as input, the last section of the overview enumerates the gear analyses that have been run successfully on any session within the project, and enumerates the runtimes for these processes. The results are visualized in a bar chart (**Figure 5**).

**Figure 5:**
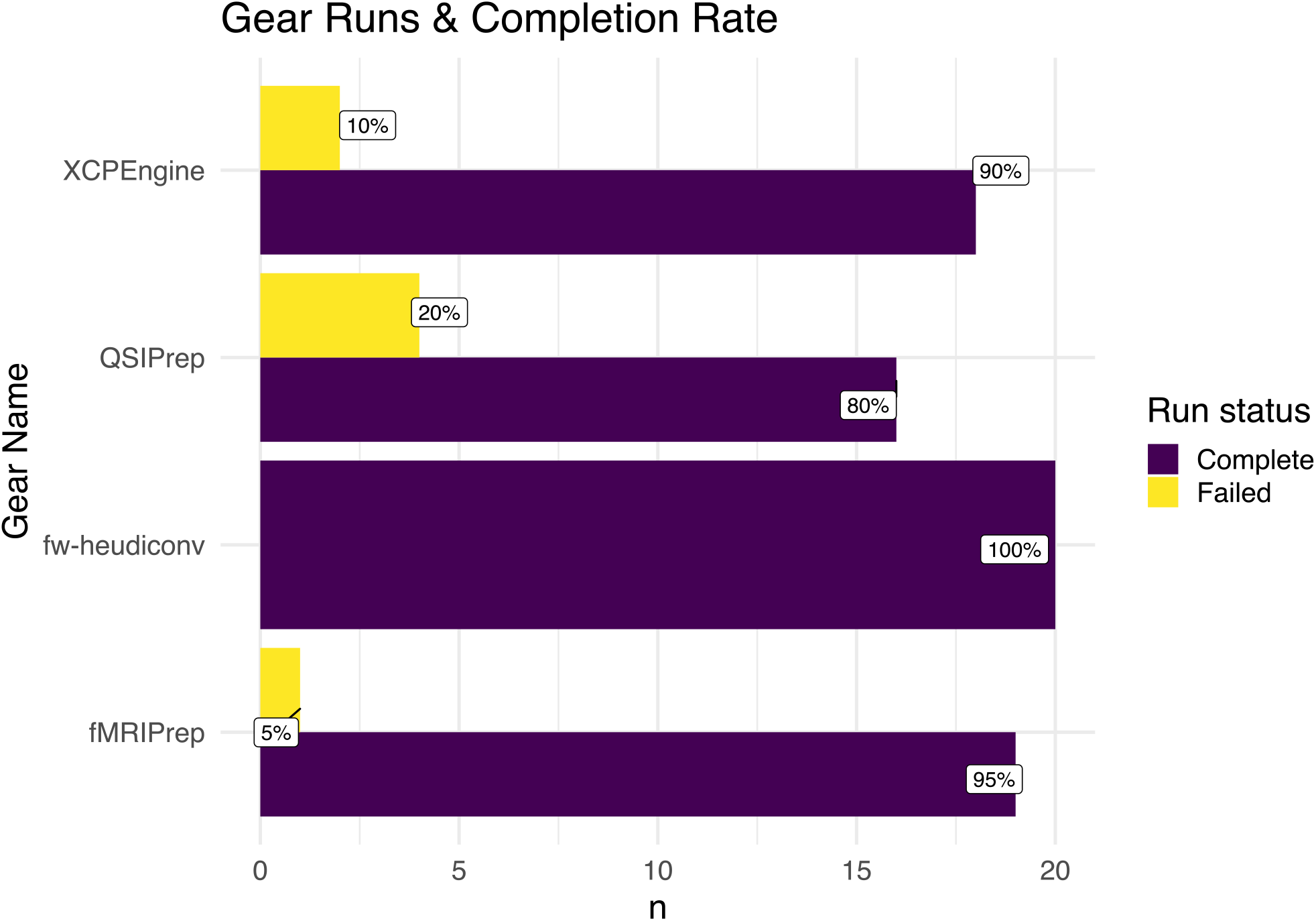
Enumeration of gear runs in a Flywheel project. The number of gear runs is shown for various gears. For each gear, the percent of completed versus failed runs is shown. For example, 95 percent of the subjects (n=19) were successfully run through fMRI-prep.

### 3.5 Flaudit: Project Completeness

As an optional input, *flaudit* allows users to specify a *template* subject — a subject from the Flywheel project who serves as an exemplar for other subjects to be compared against. This subject should be chosen based on the fact that they have both complete input data and analyses. This allows *flaudit* to determine if other subjects in the project have equally complete data or are missing specific raw data or analytic output. The project completion section of the *flaudit* report consists of three interactive tables.

In the first table, it’s assumed that the template subject has acquired a complete set of imaging sequences. These sequences are listed as columns in the table. Each subsequent row is a subject in the project, and each value in the table is a boolean (complete or incomplete) indicating if that subject has each sequence. The table is searchable, meaning that users can simply filter each column for “incomplete” to learn which subjects do not have the same data as the template (**Figure 6A**). Likewise, the second table illustrates the completeness of BIDS data for other subjects in comparison to the template (**Figure 6B**). In this case, rows indicate subjects while columns delineate the specified BIDS naming template. Lastly, the third table illustrates completeness of analytic gear runs. Researchers can use this table to compare the analytic output of all other subjects in the project to the analysis pipelines run for the template subject. To ensure uniform versions of pipeline software, the version of a pipeline that was used for each subject must match that of the template subject (**Figure 6C**).

**Figure 6:**
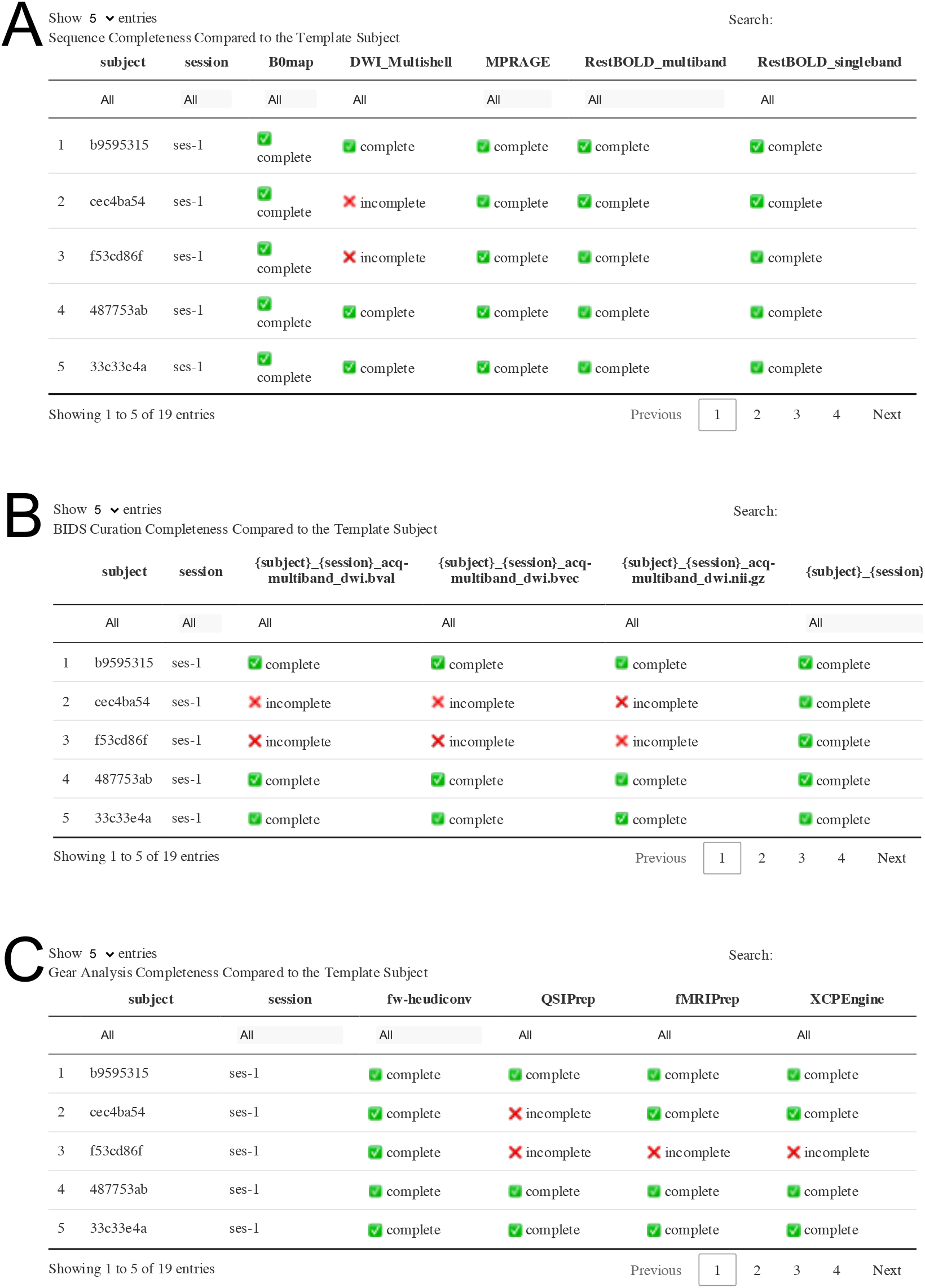
Project completeness tables compared to the template participant in a Flywheel project. Panel (**A**) compares the sequences available for each participant to the template subject and identifies missing sequences. For example, this table illustrates that subjects cec4ba54 and f53cd86f did not have DWI sequences collected. Panel (**B**) similarly shows completeness of BIDS curation. As expected, the two participants who did not have DWI sequences (in **A**) did not have diffusion data curated into BIDS. Panel (**C**) shows gear run completion; here, flaudit reports that the same two participants that lacked DWI data did not have a successful run of QSIPrep. Finally, the report notes that participant f53cd86f did not yet complete fMRIPrep or XCPEngine successfully.

## 4 DISCUSSION

FlywheelTools provides new capabilities for the popular and powerful Flywheel platform, allowing researchers to maximize reproducibility and enhance scalability. Specifically, *fw-heudiconv* provides users a flexible and reproducible way of curating data into BIDS on Flywheel. Complementary function is provided by *flaudit*, which provides intuitive visual reports of raw, curated, and processed data.

Flywheel has been rapidly adopted by major imaging centers due to its ease of use, extensive functionality, scalability, and emphasis on reproducible research. Despite these strengths, at present there have been limited options for conversion of imaging data to BIDS format on Flywheel. This step is absolutely critical, as BIDS provides a standardized format for important imaging metadata. Notably, initial curation to BIDS has frequently been an important gap in workflows for reproducible research.

Accordingly, *fw-heudiconv* provides critical functionality for reproducible and flexible BIDS curation on Flywheel. The combination of containerized code and heuristics that are version controlled with git maximizes reproducibility, ensuring that all curation steps have a clear audit trail. Furthermore, the flexible architecture employed by *fw-heudiconv* allows workflows to be updated to accommodate both new scanning protocols and the evolving specifications of the BIDS standard.

Once data are curated with *fw-heudiconv*, *flaudit* allows users to audit the data in a Flywheel project. Specifically, *flaudit* provides intuitive summaries at each stage of a typical workflow, concisely visualizing raw data, curated data in BIDS, and data processed by containerized analytic gears. These accessible reports allow users to rapidly assess the overall organizational state of a project, while interactive tables allow for more granular inspection of data. This approach facilitates understanding the diverse data types typically collected in multi-modal imaging studies.

There are of course limitations of FlywheelTools. First, it should be acknowledged that FlywheelTools is built for the Flywheel platform, and as such does not generalize to other imaging databases that are in use. However, given the rapid adoption of this platform by the imaging community, we anticipate that this toolkit will fill an important need for the many large research institutions that rely upon Flywheel. Second, some understanding of Python is necessary to build the heuristic for *fw-heudiconv*. We attempt to minimize this issue by providing both extensive documentation and heuristic templates for various uses of *fw-heudiconv*^2^, but usage is ultimately a programming task.

FlywheelTools provides essential functionality to the Flywheel platform. The flexible toolkit allows for curation and description of complex imaging studies. Taken together, the toolkit is designed to accelerate reproducible imaging research at scale.

## 6 AUTHOR CONTRIBUTIONS

T.M.T. and M.C. contributed to the design and implementation of the code and software. T.D.S., M.B., and M.C. supervised writing of the manuscript. A.A., M.C., E.R.B., D.D. and W.T. provided substantial software use-cases, software feature requests, test data, and critical bug reports. K.M. and S.L. assisted with software documentation. M.A.E., G.K.A., P.AC., J.A.D., and T.D.S. were involved in proposing, planning, and supervising all the work.

## 8 ACKNOWLEDGMENTS

None.

## Footnotes

1 Flywheel also provides a MATLAB SDK, however we use the term SDK in this work to refer to the Python SDK, which we use exclusively in FlywheelTools.

2 https://fw-heudiconv.readthedocs.io/en/latest/index.html

## REFERENCES

Banker, Kyle. 2011. MongoDB in Action. USA: Manning Publications Co.

Biehl, Matthias. 2016. RESTful Api Design. Vol. 3. API-University Press.

Book, Gregory A, Michael C Stevens, Michal Assaf, David C Glahn, and Godfrey D Pearlson. 2016. “Neuroimaging Data Sharing on the Neuroinformatics Database Platform.” Neuroimage 124: 108–992.

Botvinik-Nezer, Rotem, Felix Holzmeister, Colin F Camerer, Anna Dreber, Juergen Huber, Magnus Johannesson, Michael Kirchler, et al. 2020. “Variability in the Analysis of a Single Neuroimaging Dataset by Many Teams.” Nature, 1–7.

Cieslak, Matthew, Philip A Cook, Xiaosong He, Fang-Cheng Yeh, Thijs Dhollander, Azeez Adebimpe, Geoffrey K Aguirre, et al. 2020. “QSIPrep: An Integrative Platform for Preprocessing and Reconstructing Diffusion Mri.” bioRxiv.

Craddock, Cameron, Sharad Sikka, Brian Cheung, Ranjeet Khanuja, Satrajit S Ghosh, Chaogan Yan, Qingyang Li, et al. 2013. “Towards Automated Analysis of Connectomes: The Configurable Pipeline for the Analysis of Connectomes (c-Pac).” Front Neuroinform 42.

Esteban, Oscar, Christopher J Markiewicz, Ross W Blair, Craig A Moodie, A Ilkay Isik, Asier Erramuzpe, James D Kent, et al. 2019. “fMRIPrep: A Robust Preprocessing Pipeline for Functional Mri.” Nature Methods 16 (1): 111–16.

Halchenko, Yaroslav, Mathias Goncalves, Matteo Visconti di Oleggio Castello, Satrajit Ghosh, Michael Hanke, Matthew Brett, Taylor Salo, et al. 2018. Nipy/Heudiconv: Heudiconv V0.5.1 (version v0.5.1). Zenodo. https://doi.org/10.5281/zenodo.1306159.

Helmer, Karl G, Jose Luis Ambite, Joseph Ames, Rachana Ananthakrishnan, Gully Burns, Ann L Chervenak, Ian Foster, et al. 2011. “Enabling Collaborative Research Using the Biomedical Informatics Research Network (Birn).” Journal of the American Medical Informatics Association 18 (4): 416–22.

Herrick, Rick, William Horton, Timothy Olsen, Michael McKay, Kevin A Archie, and Daniel S Marcus. 2016. “XNAT Central: Open Sourcing Imaging Research Data.” NeuroImage 124: 1093–6.

Landis, Drew, William Courtney, Christopher Dieringer, Ross Kelly, Margaret King, Brittny Miller, Runtang Wang, Dylan Wood, Jessica A Turner, and Vince D Calhoun. 2016. “COINS Data Exchange: An Open Platform for Compiling, Curating, and Disseminating Neuroimaging Data.” NeuroImage 124: 1084–8.

Merkel, Dirk. 2014. “Docker: Lightweight Linux Containers for Consistent Development and Deployment.” Linux J. 2014 (239).

Poldrack, Russell A, and Krzysztof J Gorgolewski. 2017. “OpenfMRI: Open Sharing of Task fMRI Data.” Neuroimage 144: 259–61.

R Core Team. 2019. R: A Language and Environment for Statistical Computing. Vienna, Austria: R Foundation for Statistical Computing. https://www.R-project.org/.

Rogovin, O, Y Zhao, S Chen, Z Wang, O Papaemmanouil, SD Van Hooser, and others. 2020. “NDI: A Platform-Independent Data Interface and Database for Neuroscience Physiology and Imaging Experiments.”

Vaccarino, Anthony L, Moyez Dharsee, Stephen Strother, Don Aldridge, Stephen R Arnott, Brendan Behan, Costas Dafnas, et al. 2018. “Brain-Code: A Secure Neuroinformatics Platform for Management, Federation, Sharing and Analysis of Multi-Dimensional Neuroscience Data.” Frontiers in Neuroinformatics 12: 28.

Van Rossum, Guido, and Fred L. Drake. 2009. Python 3 Reference Manual. Scotts Valley, CA: CreateSpace.

